# scAGCI: an anchor graph-based method for cell clustering from integrated scRNA-seq and scATAC-seq data

**DOI:** 10.1101/2024.10.20.619321

**Authors:** Yao Dong, Jiaxue Zhang, Jin Shi, Yushan Hu, Xiaowen Cao, Li Xing, Yongfeng Dong, Xuekui Zhang

## Abstract

Cell clustering plays a crucial role in the analysis of single-cell multi-omics research. Despite many methods for multi-omics integrated clustering, challenges such as noise, data sparsity, and biointerpretability analysis hider effective clustering. Recent studies have demonstrated that anchor graph learning clustering in the multiview graph domain can help alleviate sparsity and high noise, reducing runtime costs. However, addressing the heterogeneity and high noise levels in multi-omics data is crucial for obtaining a more representative anchor graph. Furthermore, the existing methods often directly obtain surface information from multi-omics for clustering purposes, neglecting the mining and utilization of higher-order correlations among different features within shared information. In response to these challenges, we propose scAGCI, a cell clustering method based on anchor graphs that integrates both scRNA-seq and scATAC-seq data. Our method captures specific and shared anchor graphs representing the properties of omics data in the process of dynamic anchor unification, and mines high-order shared information to complete the omics representation. Subsequently, clustering results are obtained by integrating the specific and shared omics representation. Extensive experiments show that our method not only outperforms 13 state-of-the-art methods in terms of clustering metrics and running time, but also that the completed omics retains the biological meaning of the original omics.

## I. Introduction

SINGLE-CELL multi-omics clustering is an important part of single-cell analysis, as it typically integrates the characteristics of different omics data for cluster analysis. For instance, the characteristics of transcriptome data and proteome data are used to cluster cells together [1], [2], providing a more comprehensive understanding of cell function and characteristics. In addition, single-cell multi-omics cell clustering can facilitate the discovery of correlations and complementarities between different omics data, thereby enhancing our insight into the regulatory mechanisms governing various biological processes within cells.

Heterogeneity and high noise pose significant challenges in single-cell multi-omics clustering [3]. Each omic has its biases, for example, scRNA-seq data is affected by environmental RNA and dropout issues, while epigenomic data (such as scATAC-seq) involves sparse binary or nearly binary measurements [4]. Additionally, the number of features measured varies widely, ranging from tens or hundreds in spatial and protein data to tens of thousands in scRNA-seq data.

In order to address these issues, researchers have devoted considerable efforts. The approach to integrating different single-cell multi-omics can be divided into four main stages. Matrix decomposition and factor analysis-based methods, like SLNMF [5], MOFA+ [6]. scAI [7], Seurat v4 [8], scABC [9], Liger [10], UnionCom [11]. Encoder-based methods, like DCCA [12], scMVAE [13], scMVP [14]. Contrastive learning-based methods, such as scMCs [15]. Graph-based methods, like scMoGNN [16], GLUE [17].

The methods based on matrix decomposition and factor analysis directly obtain the shallow low-dimensional information shared among the omics, but fail to attain the nonlinear relationship between the omics [18], [19]. Therefore, the encoder method is introduced to explore the nonlinear relationship between omics. These methods work by building different encoders [20], [21], such as zero dilatation negative binomial ZINB encoders [22], VAE with probabilistic Gaussian mixture models [23], [24], etc. They iteratively learn the underlying representation in the data to reveal the distribution of different omics further, and then reconstruct the data through the decoder to maximize information retention and reduce noise. Further, the researchers implement contrastive learning [25], [26] between encoders to extract shared information from scRNA-seq and scATAC data for fusion. Single-cell multiomics data exhibit complex associations and interactions. While these encoders can reduce dimensionality to a low-dimensional space, they also result in the loss of relationships between the data. Graph learning [27]–[29] is introduced for clustering single-cell multi-omics data, enabling the capture of relationships, including similarity and connectivity between cells, to more accurately describe internal structure. However, existing methods still lack mining and utilization of shared information from multi-omics data, indicating that further refinement is needed in mining omics information.

With the continuous advancement of sequencing technology, another major challenge of single-cell multi-omics is the expanding scale of data [30]. As graph networks aggregate rich information, their computing costs are increasing exponentially. Existing clustering methods suffer from insufficient utilization of graph information and high network computing complexity, leading to increased consumption of computing resources and limitations in clustering performance.

To address all of the above issues, we develop scAGCI, an anchor graph-based method for cell clustering from integrated scRNA-seq and scATAC-seq data. Inspired by anchor learning in the multi-view domain, we propose a strategy to collaboratively optimize anchor learning and graph representation for the fused representation of single-cell multi-omics. This approach effectively extracts specific and mine high-order information shared information from multiple omics sources to represent scRNA-seq and scATAC-seq data and organically integrates this information to obtain clustering results. The main contributions are as follows:

- We take three types of single cell multi-omics’ information into account. The first one is structure information, i.e. cell-to-cell connections, which the previous works often take. The second one is feature information of each omic, for instance, gene feature in scRNA-seq data and peak feature in scATAC-seq data. The last one is their shared high-order information, like the relationship between gene and peak, which is seldom in the existing studies, but is significant for integrating single-cell multiomics. Compared with only one piece of information, this combination information more fully represents single-cell multi-omics. They are applied to construct each omic’s specific graph and their shared high-order graph.
- To optimize the specific and shared representation of single cell multi-omics, we design a dynamic anchor learning strategy. We implemented GCNs to optimize each and shared omic’s anchors and their anchor graphs by a united and bi-directional cycle. It effectively reduces the sparsity of omic data.
- We develop a hierarchical graph attention network to further mine the higher-order information of shared representation, which helps us distinguish different cell subtypes. Furthermore, we carry out a series of comparison experiments. The results illustrate our method’s outstanding performance on single cell multi-omics clustering tasks. It fully preserves the raw single-cell omic data, which reflects regulatory elements controlling gene expression.

## II. Related work

Multi-omics can be processed as a way of multi-view. In the field of multi-view analysis, it has been observed that existing multi-view clustering algorithms are primarily based on graph models, which are time-consuming and challenging to apply to large-scale datasets in practice [31]. To enhance efficiency, researchers have proposed utilizing the relationship between anchor and data to represent the relationships among all data. Currently, there are two types of anchor graph learning: heuristic sampling strategy [32]–[34] and adaptive sampling strategy [35]–[37]. Heuristic sampling strategies have been employed in the realm of single-cell omics, where researchers have introduced a scalable and efficient anchor graph clustering algorithm [38] for scRNA-seq data. The algorithm enhances the runtime and scalability of state-of-the-art consensus clustering methods while maintaining accuracy. However, this strategy often relies on K-means or random sampling, leading to clustering results that depend on initialization and may not represent the data.

We pay more attention on the adaptive anchor selection strategy. EOMSC-CA [37] applied efficient multi-view subspace clustering and consensus anchor learning. The researchers unified the anchor selection process and the graph construction process for joint optimization, and obtained a representative consensus anchor graph. In the field of multiomics, this approach is obviously not applicable. Firstly, in the learning process, this method uses the graph structure information. While in the field of single cells, features contain rich biological information that we mainly distinguish cell types. Secondly, the method learns a shared representation from different views, completely ignores the differences between different views. In the field of multi-omics, the feature expression of cells varies greatly at different molecular levels. The shared representation not only help us to get the connection across the omics, but also miss the differences of the omic graph. Fig. 1 illustrates the differences between the EOMSC-CA model and our proposed method.

**Fig. 1.**
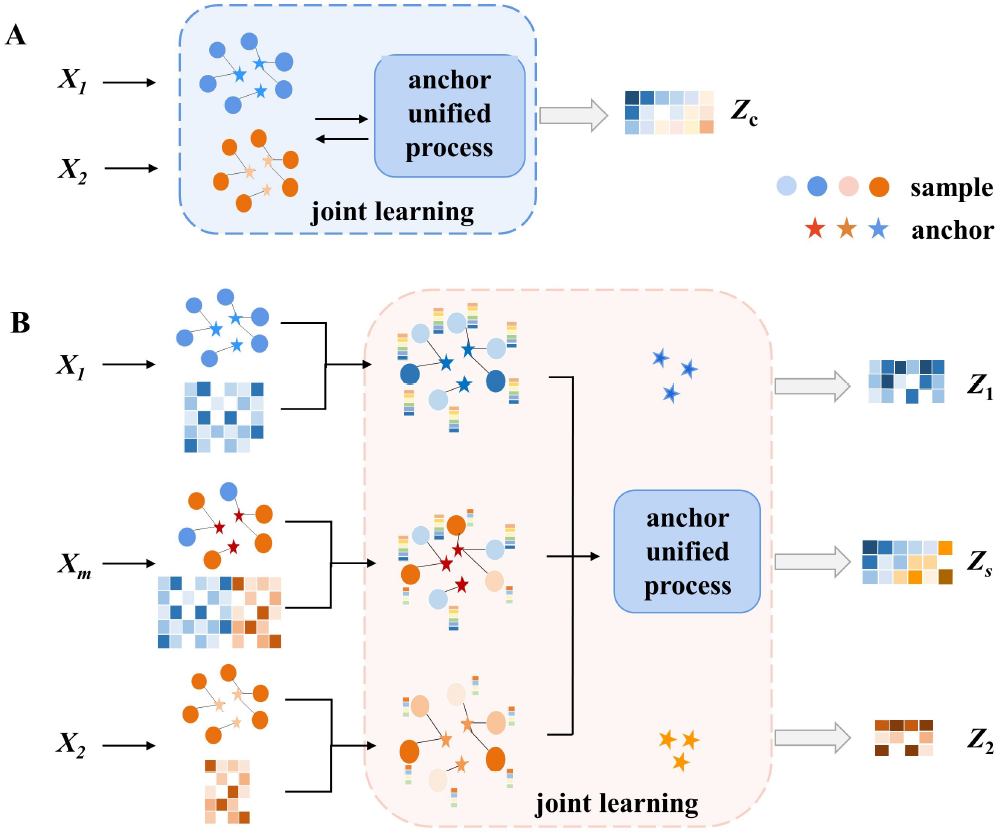
The difference on anchor graph learning mechanism between the EOMSC-CA model Fig.1A and our model Fig.1B. (i) The EOMSC-CA model only applies the structure information of graph to cluster. In contrast, we aggregate feature information of neighbors by three GCNs. (ii) By jointing learning of the anchor graph and anchor, the EOMSC-CA model learns consensus anchor ***Z***_***c***_ during the anchor unified process, while we capture not only shared representation ***Z***_***s***_ from the anchor unified process but specific information of each view ***Z***_**1**_, ***Z***_**2**_. ***X***_**1**_,***X***_**2**_ and their merging data ***X***_***m***_ are input.

## III. Methods

We develop scAGCI, an anchor graph-based method for cell clustering from integrated scRNA-seq and scATAC-seq data. It consists of three parts: multi-view subspace anchor co-optimization module, hierarchical GAT module and commonality fusion completion module. The architecture is shown in Fig. 2.

**Fig. 2.**
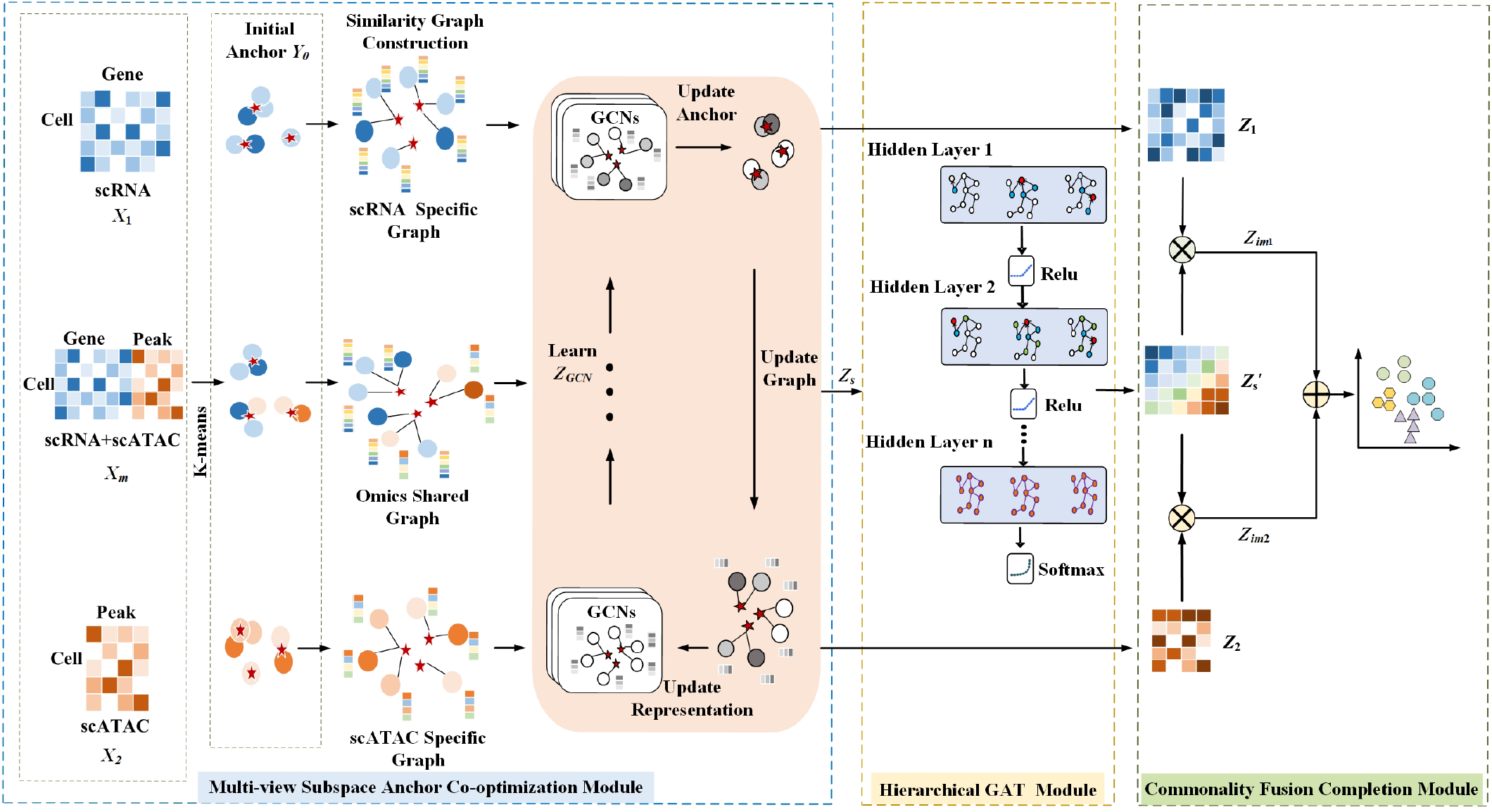
Architecture of scAGCI. Given the paired scRNA-seq ***X***_**1**_ and scATAC-seq data ***X***_**2**_ as well as their merge data ***X***_***m***_, are used as input. First of all, the multi-view subspace anchor co-optimization module uses the K-means algorithm to calculate the cluster center ***Y***_**0**_ as the initial anchor in different omics and merge omics data. We take the initial anchor and cells as nodes to construct specific and shared graphs. Next, we learn omic-specific features and structural information ***Z***_***GCN***_ by GCN, and jointly train them with dynamic anchor updates by minimizing the Frobenius norm loss to obtain omic-specific representations ***Z***_**1**_, ***Z***_**2**_, and a shared representation ***Z***_***s***_ throughout unifying the anchor. The hierarchical graph attention module is used to extract the higher-order information ***Z***^***′***^ of omics in the shared graph. Finally, we design the commonality fusion completion operator to complete the information ***Z***_***im*1**_, ***Z***_***im*2**_, and use K-means to clustering.

**Fig. 3.**
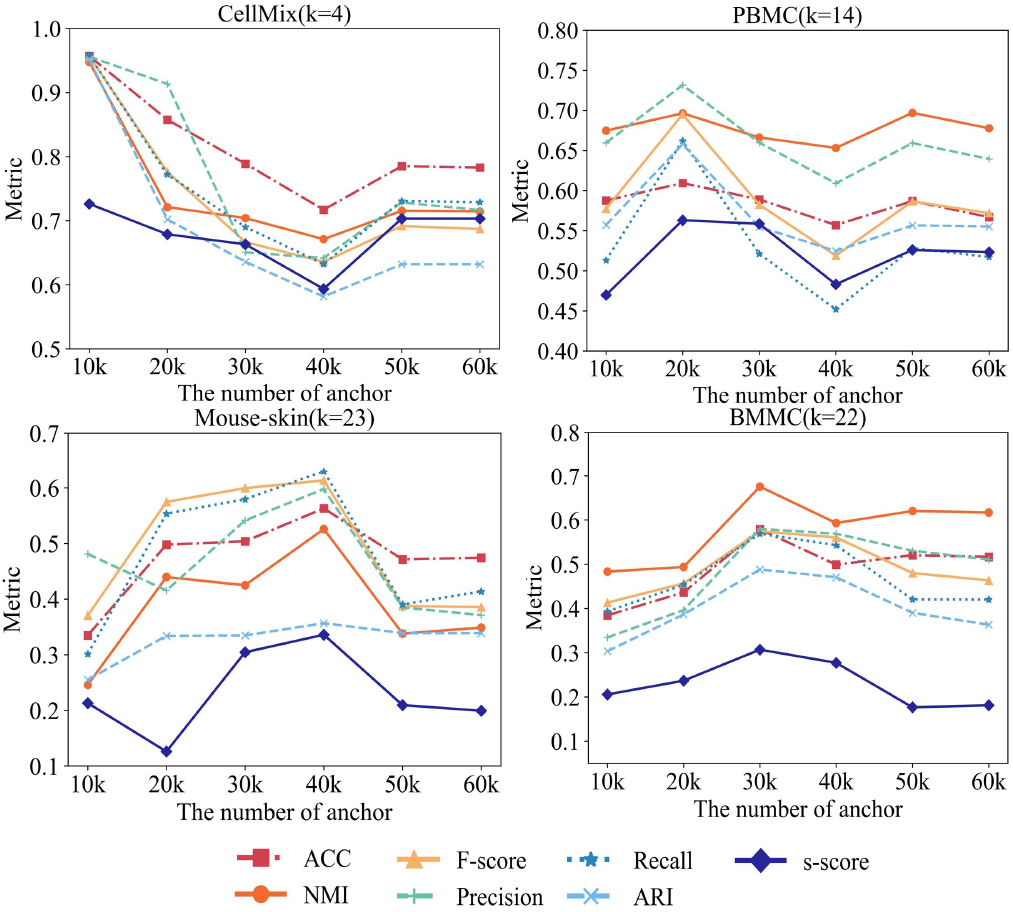
Effects of parmeter ***m*** on the 7 evaluation measures under different datasets. ***k*** denotes the number of cluster of the dataset.

### A. Multi-view Subspace Anchor Co-optimization Module

#### 1) Specific Anchor Graph Construction

With the single-cell data, the specific anchor graph is constructed in each omic.

In our study, we regard scRNA-seq data as *X*_1_, scATAC-seq as *X*_2_, and their merge data as *X*_*m*_ to form views, where 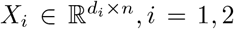 represents the *i*-th view data with *d*_*i*_ dimensions and *n* cells. 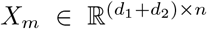 represents the merge view data with *d*_1_ + *d*_2_ dimensions and *n* cells. Assume that a linear combination of other data points in the same subspace can express each data point. The subspace clustering is to construct a graph:

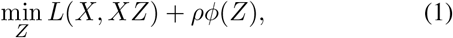

where *L*(*·*) and *ϕ*(*·*) represent the loss function and the regularization function, respectively, and *ρ* controls the trade-off between the two terms. *Z* ∈ ℝ^*n×n*^ denotes the cell similarity matrix.

Define the bipartite graph as *B*_*i*_ = (*X*_*i*_, *Y*_*i*_, *Z*_*i*_), *i* = 1, 2. The bipartite graph anchor subspace [37] clustering can be expressed as:

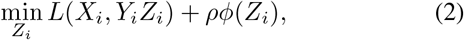

where 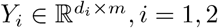 denotes the specific anchor matrix on *X*_1_ and *X*_2_. We start by randomly sampling a subset of *m* cells from all *n* cells as initial anchors *Y*_0_, which are the centroids of the identified clusters by applying k-means [39] to the above random subsample. *Z*_*i*_ ∈ ℝ^*m×n*^ is the similarity matrix between *Y*_*i*_ and *X*_*i*_. *B*_*i*_ is the anchor graph. The Frobenius norm is used as the loss function. Then, the above bipartite graph anchor subspace clustering is represented as:

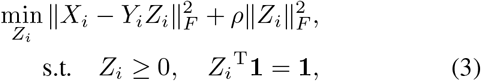

where **1** is a column vector of ones, ensuring that the sum of each column is 1. The omic-specific graph *B*_*i*_ is obtained.

#### 2) Shared Anchor Graph Construction

To achieve sufficient integration of omics information and obtain shared information between views, we define the anchor projection matrix 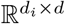 to align a shared anchor *Y*_*s*_ with the original data of the *i*-th omics. The shared graph is established among multiple omics data:

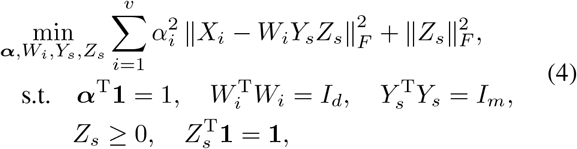

where, *v* denotes the number of views, ***α*** is the weight coefficient balancing the influence between omics and a column vector of *α*_*i*_. *X*_*i*_ is the feature matrix of the *i*-th omics, *Y*_*s*_ ∈ ℝ^*d×m*^ represents the anchor matrix on the shared anchor graph, *m* is the number of shared anchors. *Z*_*s*_ is the shared matrix between the anchor sets and omics. It also denotes the shared subspace representation. *B*_*s*_ = (*X, Y*_*s*_, *Z*_*s*_) is the shared graph.

#### 3) Anchor Selection and Graph Representation Cooperative Optimization

The existing anchor-based multi-view subspace methods largely rely on sampling strategies, resulting in a separation between anchor selection and graph representation. It contributes to the ambiguity of the final cluster structure in the constructed graph. For the challenge, we propose a cooperative optimization of graph learning representation and anchor selection to learn both the specific and shared representations of omics information. As high-dimensional data from different omics can share low-dimensional subspaces, the learned anchors in the subspace should exhibit unification.

In the optimization process, we learn each view graph representation by GCNs [40] :

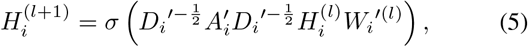

where 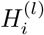 is the node representation matrix (feature matrix) of *i*-th view data at layer *l*, 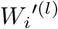 is the weight matrix at layer 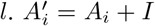 is the adjacency matrix *A*_*i*_ of *i*-th view data augmented with self-connections, forming a diagonal matrix. 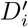 is the degree matrix of 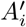. *σ* is the activation function, typically Relu. The graph convolutional operations allow nodes to propagate feature information through their neighbors, enabling the learning of node representations in the graph structure and feature matrix. We use a widely employed 2-layer GCNs, which computes the node embedding 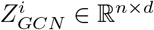 with

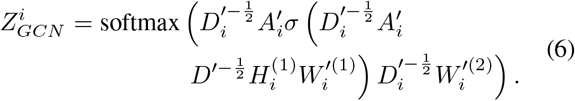

For the shared graph, during GCNs, anchor graph representation *Z*_*s*_ is equal to solve the following equation:

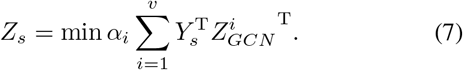

With *Z*_*s*_, *Y*_*s*_ and *α*_*i*_ being fixed, *W*_*i*_ can be optimized as

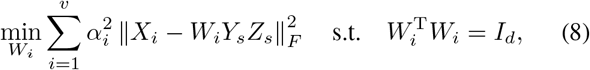

because *W*_*i*_ is independent in each view, we can express Equation (8) in the following equivalent formulation:

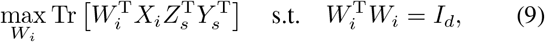

where Tr(*·*) represents the trace of a matrix. With *W*_*i*_, *Y*_*s*_ and *α*_*i*_ being fixed, *Z*_*s*_ can be optimized as

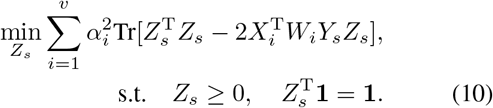

With *Z*_*s*_, *W*_*i*_ and *α*_*i*_ being fixed, *Y*_*s*_ can be written as

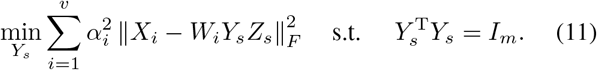

The above equation equals to the following form by unrelated section:

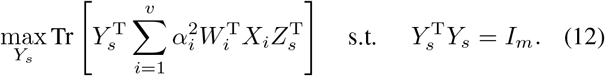

When *Z*_*s*_, *W*_*i*_ and *Y*_*s*_ are fixed, the objection function with respect to *α*_*i*_ can be formulated as

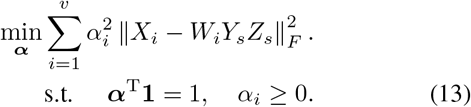

Define 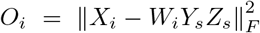, the optimal *α*_*i*_ can be obtained using the Cauchy-Buniakowsky-Schwarz [41] inequality, given by:

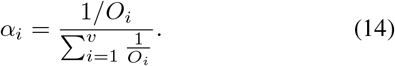

In this way, we can obtain the shared graph representation *Z*_*s*_. Using the same method, optimizing Equation 3 allows us to obtain the representation of the specific graph representation *Z*_1_, *Z*_2_.

### B. Hierarchical GAT Module

The current research emphasizes the importance of prior knowledge of single cell multi-omics (gene-gene co-regulation, peak interactions), such as in methods like sc-MoGNN and GLUE, which use prior information to construct graph networks. Our method, on the other hand, extracts shared higher-order genomics information 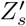 through multilayer graph attention networks to capture the interaction information between genes carried by cells. Specifically, the Hierarchical GAT [28] module employs a masked self-attention mechanism. The input *Z*_*s*_ to the graph attention layer is the node feature set *H* = {*h*_1_, *h*_2_, …, *h*_*n*_}, where *h*_*i*_ ∈ ℝ^*F*^, and the output is 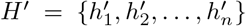, where 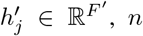 is the number of cells, 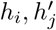 are the input and output node embedding, *F, F*^*′*^ are the dimensions of input and output node embedding. The normalized input node embedding can be represented as

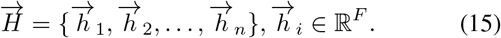

The output node embedding are

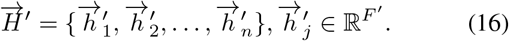

To calculate the weight of each neighboring node, a shared weight matrix **W** is applied to each node through an attention mechanism. The attention coefficient is calculated as follows:

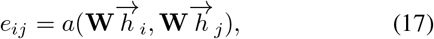

where *a* represents the importance of node *j* relative to node *i*. Normalizing the weight coefficients:

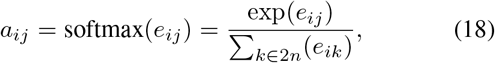

and applies the LeakyReLU, then:

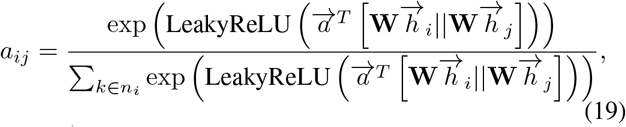

where 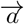 is the normalized attention coefficient calculated by *a, n*_*i*_ is the neighborhood of nodes in the graph, and || represents concatenation. The representation of node *i* is then given by:

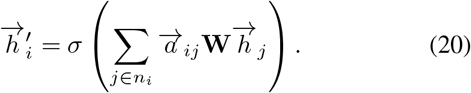

Using a multi-head attention mechanism, the final representation is obtained by taking the average:

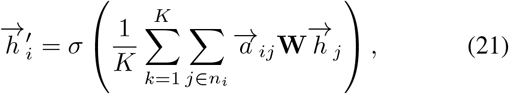

where *K* is the number of attention heads. The hierarchical GAT module updates the hidden states of all nodes in the shared graph, allowing the network to learn high-order neighboring node information, capturing hierarchical graph node information and obtaining high-order shared graph representation 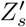.

### C. Commonality Fusion Completion Module

We design a commonality fusion completion operator to integrate information from the specific anchor graph representation *Z*_1_, *Z*_2_ and the high-order shared graph representation 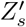, further enriching the representation of the specific graph:

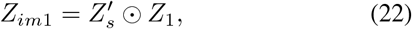

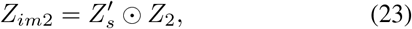

⊙ represents Hadamard product. *Z*_*im*1_, *Z*_*im*2_ represents the embedding of specific graph after imputing. Then we perform clustering by combining the representations from the specific graph and the shared graph:

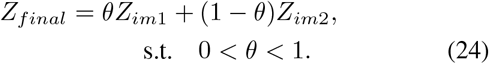

### D. Complexity Analysis

Due to the application of anchor graph learning, scAGCI has a low computational complexity analysis. It mainly involves the iterative update data of four variables: the shared or specific graph representation *Z*, the anchor matrix *Y*, the weight coefficient *α*_*i*_ and anchor projection matrix *W*_*i*_, derived from SVD, graph learning and matrix multiplication. It costs *O*(*n*^2^ + *nml* + *ml*^2^ + *m*^3^), where *m << n, l << n*. The graph attention module costs *O*(*m*^2^) and the generic fusion completion module, which mainly involves matrix multiplication and linear computation, costs *O*(*nm*).

## IV. Data and Experiments Design

### A. Datasets and Pre-Processing

In our study, we utilize three multi-omics datasets from different platforms.

1. Cell Line Mixture (CellMix) Dataset. CellMix is composed of 1047 cells with paired scRNA-seq and scATAC-seq data, which are derived from mixtures of cultured human BJ, H1, K562, and GM12878 cell lines using SNARE-seq [42]. We use the Seurat package [8] to retain the top 500 highly expressed genes in 1047 cells in the scRNA-seq dataset and the 7136 peaks within 100 kbp upstream and gene body in scATAC-seq.
2. Human Peripheral Blood Mononuclear Cell Dataset (PBMC-3K). PBMC-3K includes 3012 cells with paired scRNA-seq and scATAC-seq [43]. We first remove 200 low-quality cells (mitochondria proportion greater than 25%, threshold lower than 200). By using Azimuth software [46] to predict 2900 cell types and delete cell type data with less than five cells, obtain 2874 cells, 14 cell types. Finally, to improve efficiency, we use the Seurat package [8] to retain the top 1500 highly expressed genes in the scRNA-seq data and the 11063 peaks in the 100kbp upstream and gene bodies in the scATAC-seq data.
3. Mouse Skin Dataset (Mouse-Skin). Mouse-Skin comprises 34,774 cells with paired scRNA-seq and scATAC-seq obtained from adult mouse skin using SHARE-seq [44]. The dataset is pre-processed and we do not perform other operations on its cell characteristics.
4. Bone Marrow Mononuclear Cells Dataset (BMMC). BMMC was collected of 12 healthy human donors to support Multimodal Single-Cell Data Integration Challenge at NeurIPS 2021, 120,000 single cells from the human bone marrow with two commercially-available multi-modal technologies [45]. We input the pre-processed data into other comparison models separately. For our method, we input the two groups separately. The omics data are used as two omics views and then the data after feature merging are input into the model together as a merging view.

### B. Comparison Methods and Evaluations Metrics

Our method is compared with 13 other state-of-the-art clustering methods. Mainly include classic methods such as K-means(features-merged dataset), methods based on matrix or factor decomposition, such as Liger [10], MOFA+ [6], scAI [7], UnionCom [11], JSNMF [47], encoder-based methods DCCA [12], scMVAE [13], scMVP [14], methods based on contrastive learning, scMCs [15] and anchor graph learning-based clustering, EOMSC-CA [37]. Except for anchor graph clustering, which is a multi-view method, the others are single-cell multi-omics clustering methods. The specific parameter settings are in accordance with the original study.

We evaluate the performance of the clustering result using accuracy(ACC), normalized mutual information (NMI), F1-score, precision, recall [48], [49], adjusted rand index (ARI), Silhouette Coefficient(S-score) [50], [51]. The range of ACC, NMI, F-score, precision, Recall and ARI are all [0,1]. The S-score is [-1,1]. Some require ground truth labels and we use the cell-type labels provided referring section B. These evaluations are carried out on three data sets of different sizes and are compared with our method visually. These indicators comprehensively consider the accuracy, stability and consistency of clustering results.

### C. Experiments Design

We initialize the pre-training GCN and set it to 2 layers. The initial learning rate is 0.01, iterations are 1000 times, the weight decays weight decay set 5e-3 and the Adam optimizer is used for learning. Besides, the number of anchor needs to be determined within the range [10*k*, 20*k*, 30*k*, 40*k*, 50*k*, 60*k*], where *k* denotes the number of clusters in the dataset. For GAT, we set the number of GAT layers to 2, and the multi-head attention to 3. The completion operator set to 0.5:0.5. We conduct 10 experiments using random seeds to ensure randomness, obtaining average results for ACC, NMI, F-score, precision, recall, ARI, and S-score, while validating the results every 30 iterations.

As for the comparison methods, we adopt the parameters that yielded the best results in the original paper. All algorithms are run on a machine with Intel(R) Xeon(R) Gold 6348 CPU @ 2.60GHz addressing 100GB memory and one NVIDIA A30 TENSOR CORE GPU. scAI, EOMSC-CA and JSNMF are implemented on MATLAB(version R2021a). Both MOFA+ and Liger are implemented on Rstudio(version 2021). Multi-omics integration methods should fully retain the specific biological [52], [53] information of single omics datasets and produce embedding with good cluster discrimination after integration. To further comprehensively evaluate the performance of our method, we apply 7 indicators to compare with 13 algorithms and make the visualization. Additionally, we perform differential gene expression and enrichment analyses to identify genes with significantly altered expression under different conditions, further exploring their functions. Moreover, we analyze the transcription factor motifs for each cell to investigate their role in gene regulation. Finally, we examine the runtime of 4 categories of methods.

## V. Results and Discussion

### A. Evaluate algorithm performance in comparison with SOTA methods

The table II shows the clustering results of our method and 13 other methods. The values marked in bold are the best results among all methods, and the values underlined are suboptimal. We conduct the following findings. We observe that scAGCI always achieves satisfactory embeddings compared with other methods. For small to medium-sized datasets, we find that compared to the suboptimal method scMCs, our method has largely improved, with indicators improved by 1.32%, 4.06%, 1.83%, 2.32%, 1.35%, 1.31% and 5.99% on ACC, NMI, F1-score, Precision, Recall, ARI, S-score on the Cellmix dataset. On the PBMC-3k dataset, our method improved the index by 5.76%, 8.51%, 2.59%, 4.01%, 1.23%, 2.42% and 0.45% compared with the scMVAE-NN.

**TABLE I.** Summary of the Datasets after Pae-processing.

| Datasets | Cells | Genes | Peaks | Subtypes |
| --- | --- | --- | --- | --- |
| Cellmix | 1047 | 500 | 7136 | 4 |
| PBMC-3k | 2847 | 1500 | 11063 | 14 |
| Mouse-skin | 34774 | 2000 | 35943 | 23 |
| BMMC | 69249 | 23296 | 120010 | 22 |

**TABLE II.**
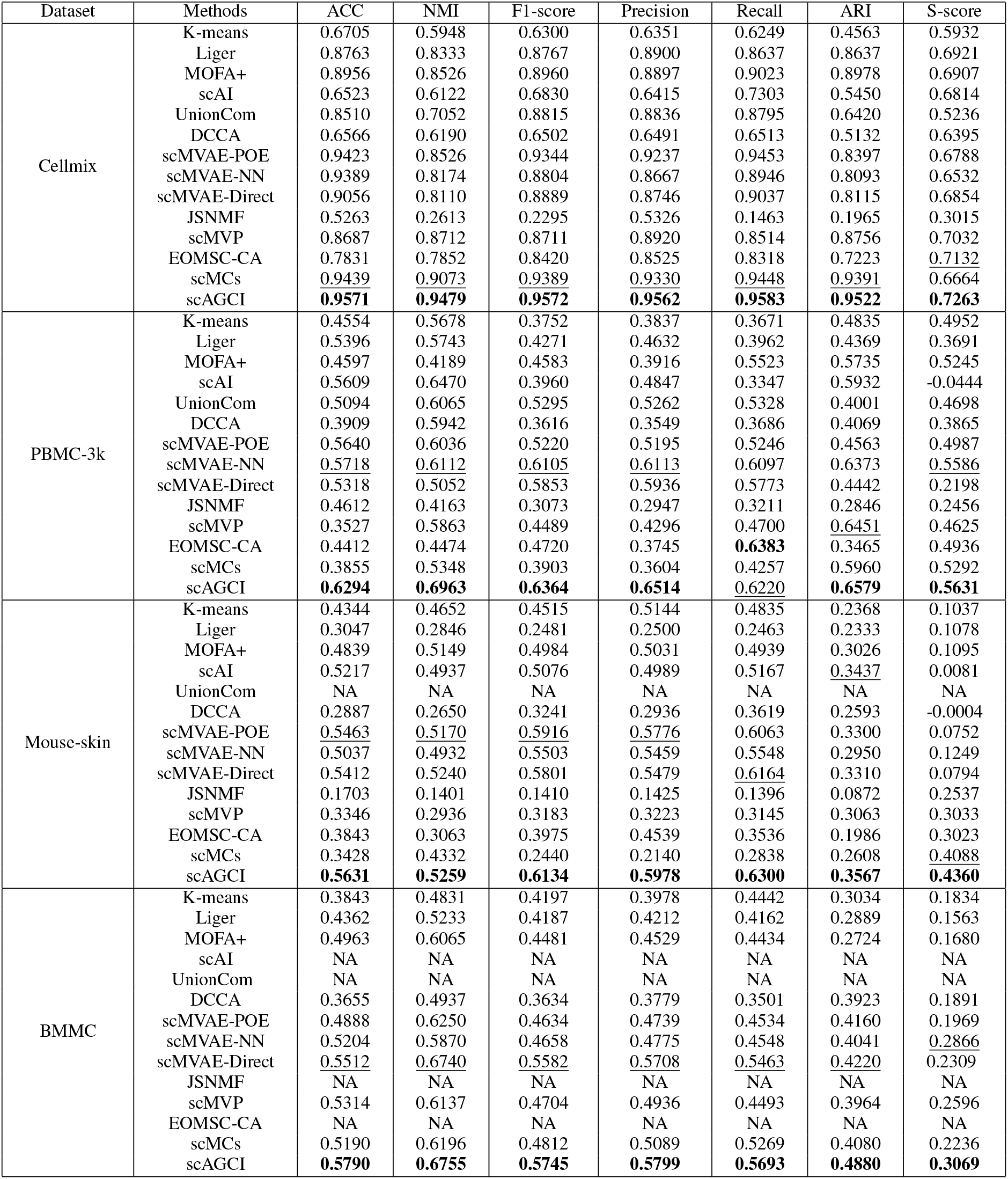
COMPARING RESULTS UNDER DIFFERENT METHODS.

For large-sized dataset, our method also has an improvement effect visible. Compared with the suboptimal method scMVAE-POE, our method has improved by 1.68%, 0.89%, 2.18%, 2.02%, 2.37%, 2.67% and 36.72%, respectively on Mouse-skin dataset. Specially, on the larger sized dataset with more than 100,000 single cells, our method has also achieved improvement of 2.78%, 0.15%, 1.63%, 0.91%, 2.30%, 6.60%, 2.03% respectively, compared to the suboptimal method scMVAE.

Its outstanding performance mainly benefits from the extraction of specific omics information, the mining of high-order shared information and the full fusion of these information. Regarding the other comparison methods, scMCs takes into account shared and specific information extraction, but it misses the high-order shared information. Other methods pay more attention to the extraction and completion of specific information from omics.

We further find that scAGCI significantly alleviates sparsity. For instance, on CellMix dataset, the matrix decomposition-based method scAI, Unioncom have ARIs of 0.5450 and 0.6420, respectively. Methods using ZINB decoders to reconstruct data like scMVAE and scMCs achieve ARI values above 0.8, while we attain an ARI of 0.9522. The lower ARI indicates that they performed poorly in addressing the sparsity of single-cell datasets. scAGCI retains better sparsity alleviation effects through the multi-view subspace anchor co-optimization method, further improving the clustering discriminability of the integrated embeddings. The results are also notable on PBMC-3k, Mouse-skin and BMMC datasets.

### B. Analysis of the Effects of Parameters on the Clustering Performance

To analyze the impact of the number of anchors on performance, we conducted experiments on four datasets, as depicted in Fig. 6. We varied the number of anchors within the range of 10k to 60k (with step size of 10k) and implemented our method on different datasets.

We have observed that the selection of anchor points needs to be adjusted based on the scale of the dataset. This suggests that utilizing anchor point clustering can greatly improve the clustering performance of single-cell multi-omics data. For the Cellmix dataset, we recommend setting the anchor point count at 10k. This allows for effective capture of key cell type features while minimizing computational overhead, given its relatively small size. For the PBMC-3k dataset, we suggest setting the anchor point count at 20k. This will enhance clustering accuracy by increasing the ability to capture the heterogeneity and subtle distinctions between various immune cell types. In the Mouse-skin dataset, we set the anchor point count at 40k, which yielded optimal results. This is due to the presence of many cells with similar gene expression profiles, enabling better differentiation among closely related cell types. For the BMMC dataset, we recommend selecting 30k anchor points. The complexity and diversity of this dataset benefit from a greater number of anchor points to represent the cellular landscape and identify unique subpopulations. Furthermore, after the number of anchor points reaches a certain value, it tends to stabilize or even decrease the effect of clustering. This indicates that the noise caused by too many cells will affect the resolution of the cells. This demonstrates that the anchor method can effectively improve the noise of the data.

Overall, For datasets with varying sizes and resolutions, adjustments to the number of anchor points are necessary.

### C. Ablation study

scAGCI consists of three main modules: the Multi-view Subspace Anchor Co-optimization Module(MAC), the Hierarchical GAT Module(H-GAT), and the Commonality Fusion Completion Module(CFC). The Multi-view Subspace Anchor Co-optimization Module extracts specific and shared information from omics data, the Hierarchical GAT Module explores high-oder shared information and the Commonality Fusion Completion Module complements specific information and integrates shared and specific information. The three modules together contribute to the excellent performance of scAGCI in integrating omics information(Table III). We conduct ablation experiments on three datasets to demonstrate the effectiveness of each module in our method.

**TABLE III.**
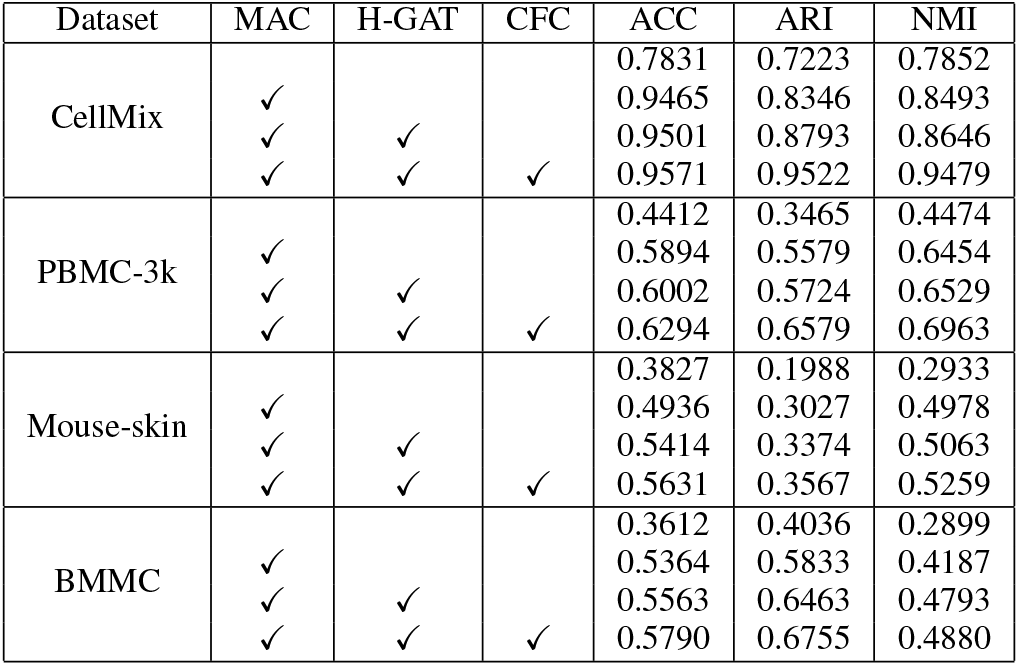
ABLATION STUDY.

Thanks to MAC module, our results are improved by 16.34%, 11.23%, 6.41% on CellMix, 14.82%, 21.14%, 19.8% on PBMC-3k, 11.09%, 10.39%, 20.45% on Mouse-skin, and 17.52%, 17.97%, 12.88% on BMMC. The results show that the specific graph representation, shared graph representation and anchor co-optimization trained by GCN are beneficial in enhancing model performance. This is mainly due to aggregating feature information such as genes and peaks into nodes to obtain more representative anchors.

On this basis, we add the H-GAT module. We observe that the clustering effect is improved by 0.36%, 4.47%, 1.53% on Cellmix, 1.08%, 1.49%, 0.75% on PBMC-3k, 4.78%, 3.47%, 0.85% on Mouse-skin and 1.99%, 6.30%, 6.06% on BMMC. The module explores the high-order information shared between the peak and gene, adaptively adjust weights for different neighbours, thus improving the model effect.

Finally, we add the CFC module and our results improves by an average of 2.02%, 5.17%, and 3.88% on 4 datasets. This illustrates that using high-order shared information to complete specific information and then fuse it helps to obtain a more differentiated cell representation while retaining the shared information of the cell.

The above ablation results demonstrate the effectiveness of each module. They also show that the complete model using all of them has better clustering results.

### D. scAGCI can distinguish cells compared to single omic

We visualize the results of single omic to demonstrate the advantages of multi-omics in identifying cell types in Fig. 4. On the CellMix dataset, raw scRNA-seq data could clearly cluster most cells into four cell clusters, but some mixing are observed in the BJ, GM, and H1 cells. The raw scATAC-seq data show a higher clustering degree for the K562 cell than others in Fig. 4A scRNA-seq, scATAC-seq. On the PBMC-3k dataset and Mouse-skin dataset, although the clustering boundaries are unclear, the clustering effect of raw scRNA-seq can effectively group similar cells together shown in Fig. 4B, C, D scRNA-seq, scATAC-seq. However, in scATAC-seq, direct clustering cannot distinguish cell types, and scATAC-seq only shows clear distinctiveness in some cell types. Compared to the clustering results of single-omics data, our integration method scAGCI effectively merges cell clusters into one class and is more discriminative in Fig. 4B, C, D.

**Fig. 4.**
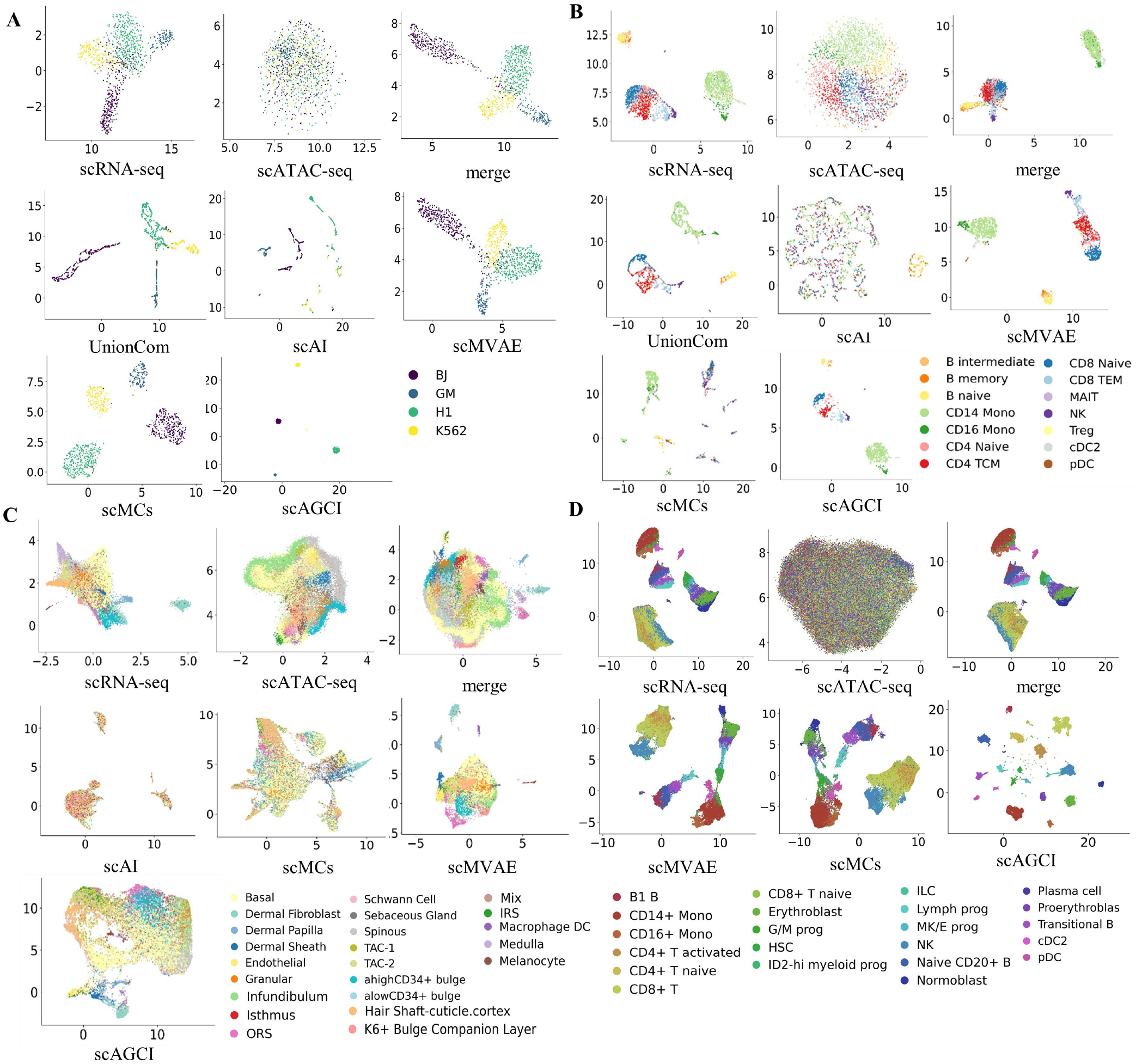
The UMAP scatterplot visualizations on CellMix, PBMC-3k, Mouse-skin and BMMC are respectively shown in subfigure A, B, C and D. Each cell is colored with their true cell type. The first row of each subfigure is the raw data input scRNA-seq, scATAC-seq, feature-merging scRNA-seq and scATAC-seq, respectively, and the visualization of our method. The other line is the visualization effect of integration methods UnionCom, scAI, scMVAE and scMCs. Some of these methods have an overflow of memory on the Mouse-skin and BMMC datasets.

### E. scAGCI further distinguishes cell subtypes

To exhibit the unique advantages of scAGCI in cell type clustering, we visualize UnionCom, scAI, scMVAE, scMCs, scAGCI and features-merging using UMAP visualization in Fig. 4.

Among the visualized integration methods on the CellMix in Fig. 4A, scAI clearly distinguishes between GM and BJ cells, but H1 and K562 cells are mixed. UnionCom, scMVAE and scMCs all differentiate the four cell types with distant inter-class distances. In contrast, our method, benefiting from the multi-view space anchor co-optimization module, tightly clusters each cell type together with smaller inter-class distances. As shown in Fig. 4B, all visualized integration methods separate B cells from other immune cells on the PBMC-3k dataset. Methods like scMVAE or UnionCom demonstrate significant inter-class distances among B cells, CD14 Mono, CD16 Mono, cDC, and various T cells, yet they struggle to accurately cluster specific subtypes like CD4 Naive, CD4 TCM, CD8 Naive, and CD8 TCM. Our method, however, clearly delineates these subtypes, showcasing its superior capability. We visualize these methods on the mouse-skin dataset as well. Unfortunately, the UnionCom method fail to run due to the dataset’s large size. Compared to scAI, other methods are able to cluster cell types such as Dermal Fibroblast, Dermal Papilla, Dermal Sheath, Macrophage DC, and Melanocyte. Ours also clusters the Schwann Cell cluster and the Hair Shaft-cuticle cluster cortex cell cluster. In Fig. 4D, scAGCI successfully differentiates between CD14+ mono and CD16+ mono, as well as activated CD4+ T cells and naive CD4+ T cells, further highlighting its effectiveness in large-scale data analysis on the BMMC dataset.

For some datasets that are not inherently differentiated by single-omics, our method can only distinguish between a small number of cell types (e.g. Mouse-skin datasets). Overall, scAGCI makes cell subtypes more distinguishable.

### F. scAGCI retains the specific information of scRNA-seq

scRNA-seq captures transcriptional profiles to reveal gene expression patterns. We take the human datasets (CellMix and PBMC-3k) as examples to verify the effectiveness of the scRNA-seq and scATAC-seq data specific information extracted by our method, we conduct differential gene expression analysis and enrichment analysis on the specific information extracted from the scRNA-seq data on the CellMix and PBMC-3k datasets. In this study, we use the Seurat package to identify the top 3 genes with the greatest differences between different cell clusters as candidate genes, then display the distribution of these differentially expressed genes in the cell clusters in Fig. 5A, B. According to known information [54], [55], the CellMix dataset consists of human embryonic stem cells (H1), B lymphocytes (GM12878), blood cancer lymphocytes (K562) and human skin fibroblasts (BJ). The PRAME, COL1A and HLA-DRB1 are considered known marker genes for K562, BJ and GM, respectively. According to the CellMarker [56] database, TERF1, ESRG and GRID2 are marker genes for embryonic tissue. Fig. 5A shows the expression of these genes in H1 cells, which is consistent with the actual differential gene expression in cells.

**Fig. 5.**
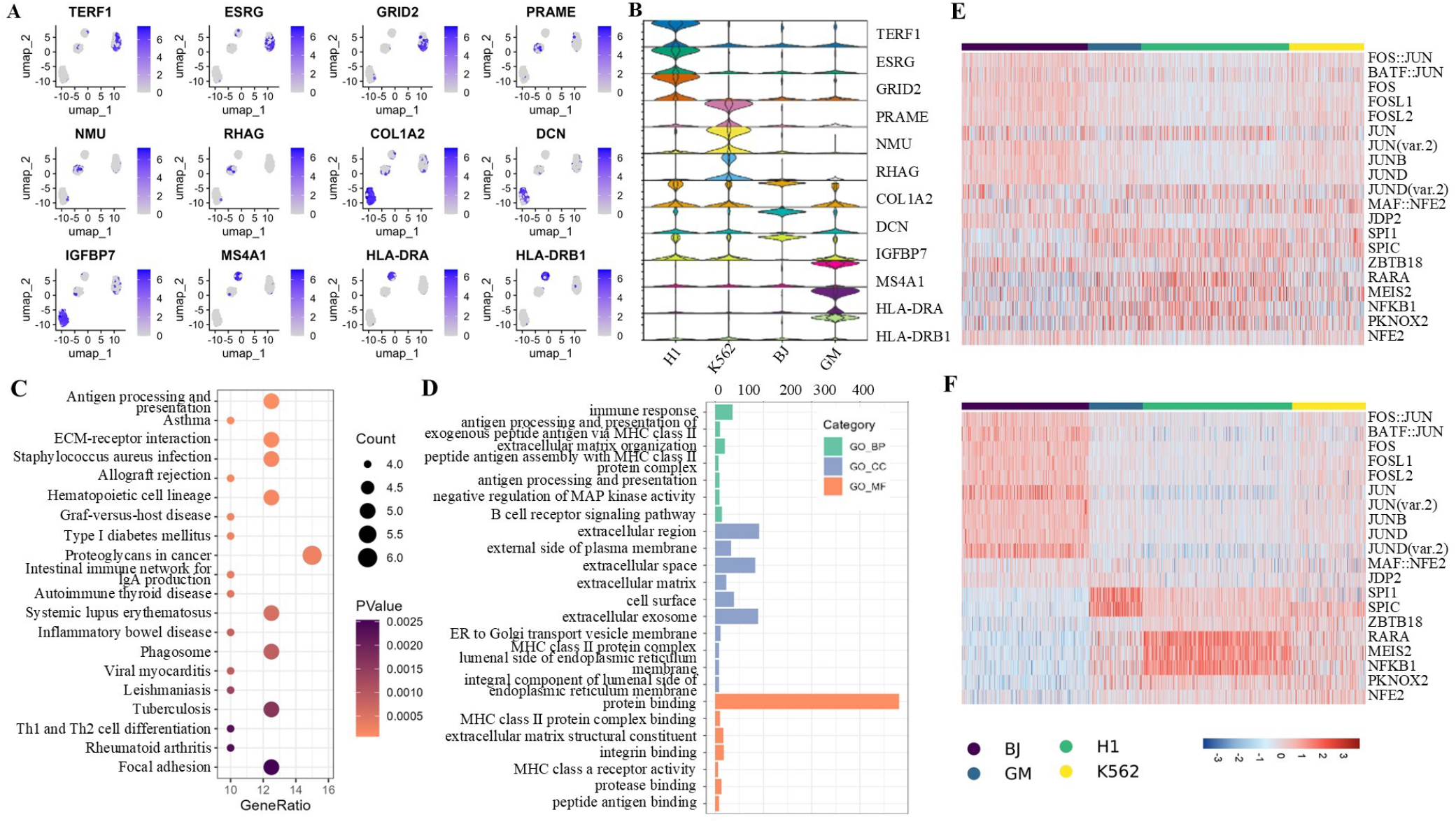
scRNA-seq and scATAC-seq analysis on Cellmix dataset. Fig. 5A shows the expression of top3 marker genes across different cell clusters. Fig. 5B shows the distribution of these top3 marker genes. Fig. 5C The bubble plot visualizes the results of enrichment analysis for KEGG pathways, which shows the ratio of the number of genes in the gene set that overlap with the genes in the pathway to the total number of genes in the pathway. Fig. 5D shows the Go enrichment. For raw data (Fig. 5E) and the extracted scATAC-seq data (Fig. 5F) transcription factor score on CellMix dataset by ChromVAR.

In addition, we perform KEGG enrichment analyses and gene ontology (GO) term enrichment on genes. In KEGG pathway analysis in Fig. 5C, we notice the enrichment in Antigen processing and presentation and Focal adhesion. The GO enrichment analysis on Cellmix data reveals several findings in Fig. 5D. We observe enrichment in molecular functions associated with protein binding, including MHC class II proteins and integrins. Biological processes related to antigen processing and presentation, antigen receptor activity are enriched. In terms of cellular components, we observe enrichment in extracellular exosomes and endoplasmic reticulum membranes.

Similarly, in the extracted scRNA-seq specific information on the PBMC-3k dataset, we select the top 2 genes with the greatest differences between each cell cluster and other cells as candidate genes in Fig. 6A, B. The PBMC-3k dataset consists of single-cell from human peripheral blood, mainly including lymphocytes (T cells, B cells and NK cells), monocytes, dendritic cells, etc. The omics-specific information of scRNA-seq blurs the boundaries among CD4 TCM and Treg cells, B intermediate cells and B memory cells, which also reflecting in previous single-omics visualizations.

In Fig. 6C, we observe primary immunodeficiency and T cell receptor signalling pathways may correspond with the immune response, inflammatory response we mentioned in BP. Besides, Hematopoietic cell lineage, Cell adhesion molecules, and Antigen processing and presentation were enriched in KEGG. In Fig. 6D, we observe cellular components term enriched in plasma membrane, integral components of membranes, and receptor complexes. In biological processes, significant enrichments are detected in signal transduction and cell surface receptor signalling. Additionally, immune response, inflammatory response, and cytoplasmic translation are also observed to be enriched. Regarding molecular functions, we view enrichments in protein binding and structural constituents of ribosomes. These findings collectively suggest that genes within this dataset play pivotal roles in cellular membrane and receptor-related activities and immune responses.

In summary, we confirm that the extracted scRNA-seq specific information can essentially recover the intrinsic information of the omics itself. These extracted omics information have clear biological bases, which helps to more accurately cluster cells and deepen the understanding of biological differences between cells.

**Fig. 6.**
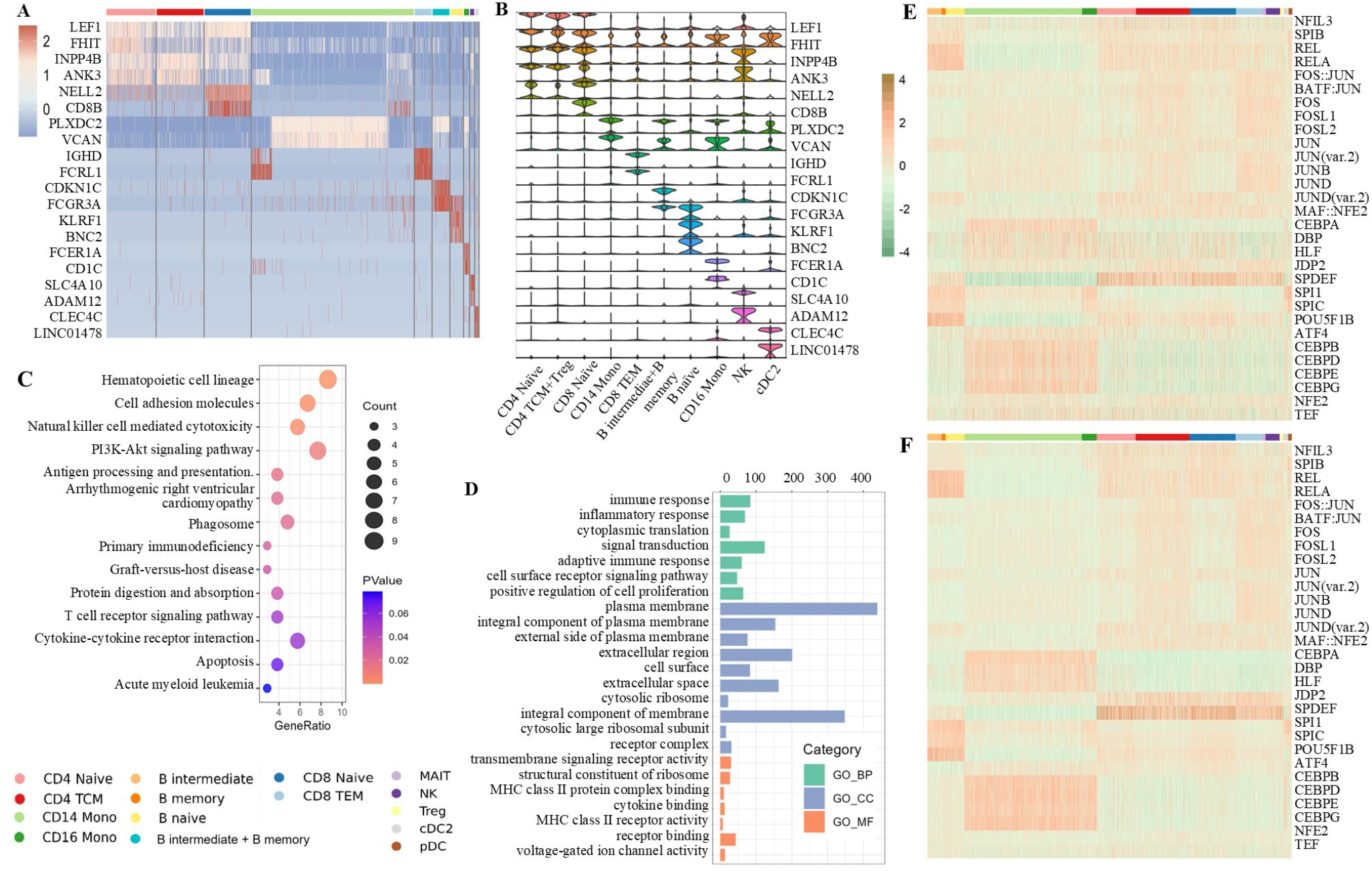
scRNA-seq and scATAC-seq analysis on PBMC-3k dataset. Fig. 6A shows the expression of top2 marker genes across different cell clusters. Fig. 6B shows the distribution of top2 marker gene expression across different cell types. Fig. 6C The KEGG bubble plot visualizes the results of enrichment analysis for KEGG pathways, which shows the ratio of the number of genes in the gene set that overlap with the genes in the pathway to the total number of genes in the pathway. Fig. 6D shows the Go enrichment. For raw data (Fig. 6E) and the extracted scATAC-seq data (Fig. 6F) transcription factor score on PBMC-3k dataset by ChromVAR.

### G. scAGCI retains the specific information of scATAC-seq

scATAC-seq elucidates chromatin region accessibility, reflecting regulatory elements controlling gene expression. In order to further illustrate the bio-interpretability and cell discrimination of our method, we calculate the TF activity score [57] for both the raw and extracted scATAC-seq specific omics data in Fig. 5E, F. It can be seen that the extracted specificity, such as JUN, JUND, FOS, BATF::JUN in BJ cells, NFK1, RARA in H1 cells, SPl1 in GM12878 cells and FOS, JUND co-associate in K562 but not in GM12878 indicates that our model effectively retains specific transcription factors between different cell types [56], [58].

We visualizes the ATAC-seq extracted from the PBMC-3k dataset in Fig. 6E, F. Although the effect is not obvious compared with CellMix, the PBMC-3k dataset has a relatively medium level of noise, while the CellMix dataset has a relatively high level of noise. However, we obverse that the original data of PBMC-3k roughly divide 14 kinds of cells into three types, three kinds of B cells, memory cells and T cells, etc. cDC2 and pDC cells are obviously different from other cells in transcription factors SPI1 and SPIC, and they are far away from other cells in other clustering methods. On the other hand, B cells have higher activity scores in REL, RELA and POU5F1B transcription factors, while the activity level of memory cells in SPDEF transcription factors is much lower than that of other kinds of cells. The expression of CD16 Mono on SPl1 and SPlB is slightly higher than that of CD14 Mono. Finally, in many T cells, the transcription factor SPDEF is highly expressed, and Naive cells have lower scores on FOS and JUN than TCM cells [59], [60]. Therefore, our method further distinguishs CD14 Mono, CD16 Mono, CD4 Naive, CD4 TCM, CD8 Naive and CD8 TCM cells.

### H. scAGCI saves time cost

Considering the increasing scale of single-cell sequencing datasets, the runtime on analysis is a critical concern. We record the time consumption of each method on datasets of varying sizes. As depicted in the Table IV, on the CellMix dataset, UnionCom has the shortest runtime, followed by our method, while scAI consumes the most time. Our method demonstrates the shortest runtime, followed by the Union-Com algorithm on the PBMC-3k dataset. On the Mouse-skin dataset, UnionCom fails to run on large-scale datasets, while our algorithm not only runs but also has the shortest runtime. In small datasets (Cellmix), the runtime of Union-Com based on matrix decomposition is more advantageous, but our method performs better in clustering effect. For medium(PBMC-3k) and large datasets (Mouse-skin, BMMC datasets), our method has obvious advantage on runtime and clustering effect.

**TABLE IV.**
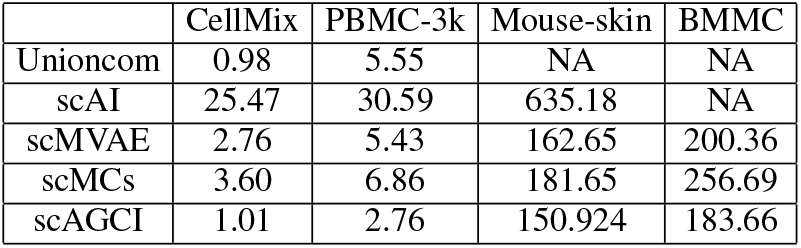
DIFFERENT SCALE DATASETS RUNTIME (MINUTE).

|  | CellMix | PBMC-3k | Mouse-skin | BMMC |
| --- | --- | --- | --- | --- |
| Unioncom | 0.98 | 5.55 | NA | NA |
| scAI | 25.47 | 30.59 | 635.18 | NA |
| scMVAE | 2.76 | 5.43 | 162.65 | 200.36 |
| scMCs | 3.60 | 6.86 | 181.65 | 256.69 |
| scAGCI | 1.01 | 2.76 | 150.924 | 183.66 |

## VI. Conclusion

In this work, we propose scAGCI, an anchor graph-based methods for cell clustering from scRNA-seq and scATAC-seq to address the challenge of heterogeneity and high sparsity, we take the structure and feature information from each omics and shared anchor graph during the dynamic anchor learning strategy. We mine the high-order shared information to rich the relationship between peak and gene. Experimental results demonstrate that scAGCI exhibits excellent capabilities in distinguishing different cell types and subtypes. scAGCI addresses the limitations of anchor graphs in representing omics data by co-optimizing anchor graph and graph embedding methods, resulting in reduced integration time for large-scale datasets, decreased sparsity and noise in omics data. While scAGCI demonstrates excellence in distinguishing cell types and subtypes, determining the optimal number of anchors poses challenges.

Multi-omics integration study helps uncover various aspects of cell function, expression dynamics, and metabolic status, leading to a deeper understanding of cell diversity and complexity. It aids in exploring disease-specific omics features and molecular mechanisms, offering insights into disease causes and mechanisms.

Future work will focus on developing more intelligent anchor selection algorithms to significantly mitigate the current limitation of requiring manual selection of the number of anchors.

## Acknowledgment

The authors thank the researchers who provide us with source code for a comparison.

## Available

The codes and supplementary document are available at https://github.com/hebutdy/scAGCI.

